# Temperature and development drive variation in oral morphology among tailed frog (*Ascaphus* spp.) populations

**DOI:** 10.1101/2020.04.06.027664

**Authors:** AS Cicchino, CM Martinez, WC Funk, BR Forester

## Abstract

Morphological variation is often maintained by complex and interrelated factors, complicating the identification of underlying drivers. Tadpole oral morphology is one such trait that may be driven by the independent and interacting effects of the environment and variation in developmental processes. Although many studies have investigated tadpole oral morphological diversity among species, few have sought to understand the drivers that underlie intraspecific variation. In this study, we investigated potential drivers of labial tooth number variation among populations of two species of tailed frogs, the Rocky Mountain tailed frog (*Ascaphus montanus*) and the Coastal tailed frog (*A. truei*). We counted labial teeth from 240 tadpoles collected across elevation (both species) and latitudinal (*A. truei*) gradients, providing a natural temperature gradient. We tested the effects of developmental stage and local temperature conditions on labial tooth number. We found that labial tooth number variation was independently affected by both developmental stage and local temperature, as well as the interacting effects of these two variables (pseudo-R^2^ = 67-77%). Our results also uncovered consistent patterns in labial tooth row formula across the ranges of both species; however, *A. truei* tadpoles from northern British Columbia had a unique bifurcation of a posterior tooth row. This study highlights the diversity in intraspecific tadpole oral morphology and the interacting processes that drive it.

## Introduction

Anuran (frogs and toads) tadpoles are a phenotypically diverse group of organisms with complex oral morphologies. Generally, the tadpole oral apparatus consists of an oral disc with papillae and multiple rows of keratinized labial teeth (*sensu* Altig and McDiarmid 1999) that surround a jaw sheath and mouth (Figure 1). These mouthparts play a critical role in feeding by stabilizing the tadpole and lifting food off of substrates (Wassersug and Yamashita, 2001; Venesky et al., 2010a; 2013). Tadpole metamorphosis is energetically costly, and tadpoles nearing and undergoing metamorphosis often have a complete re-organization of the gut and mouth which may inhibit feeding or digestion (Etkin, 1963; Metter, 1964; Toloza and Diamond, 1990). Therefore, tadpoles must acquire sufficient energy stores prior to metamorphosis (Bennett and Marshall, 2005; Orlofske et al., 2009). Because oral morphology can facilitate or constrain feeding on certain substrates (Annibale et al., 2019; 2020), the environment may select for certain morphologies, as has been suggested by the similarity of oral morphologies within some ecological guilds (Altig and McDiarmid, 1999; Altig, 2006; Vera Candioti and Altig, 2010). Oral morphology that conforms with the physical properties of the local environment is therefore likely to be important in ensuring that tadpoles survive and metamorphose.

**Figure 1.**
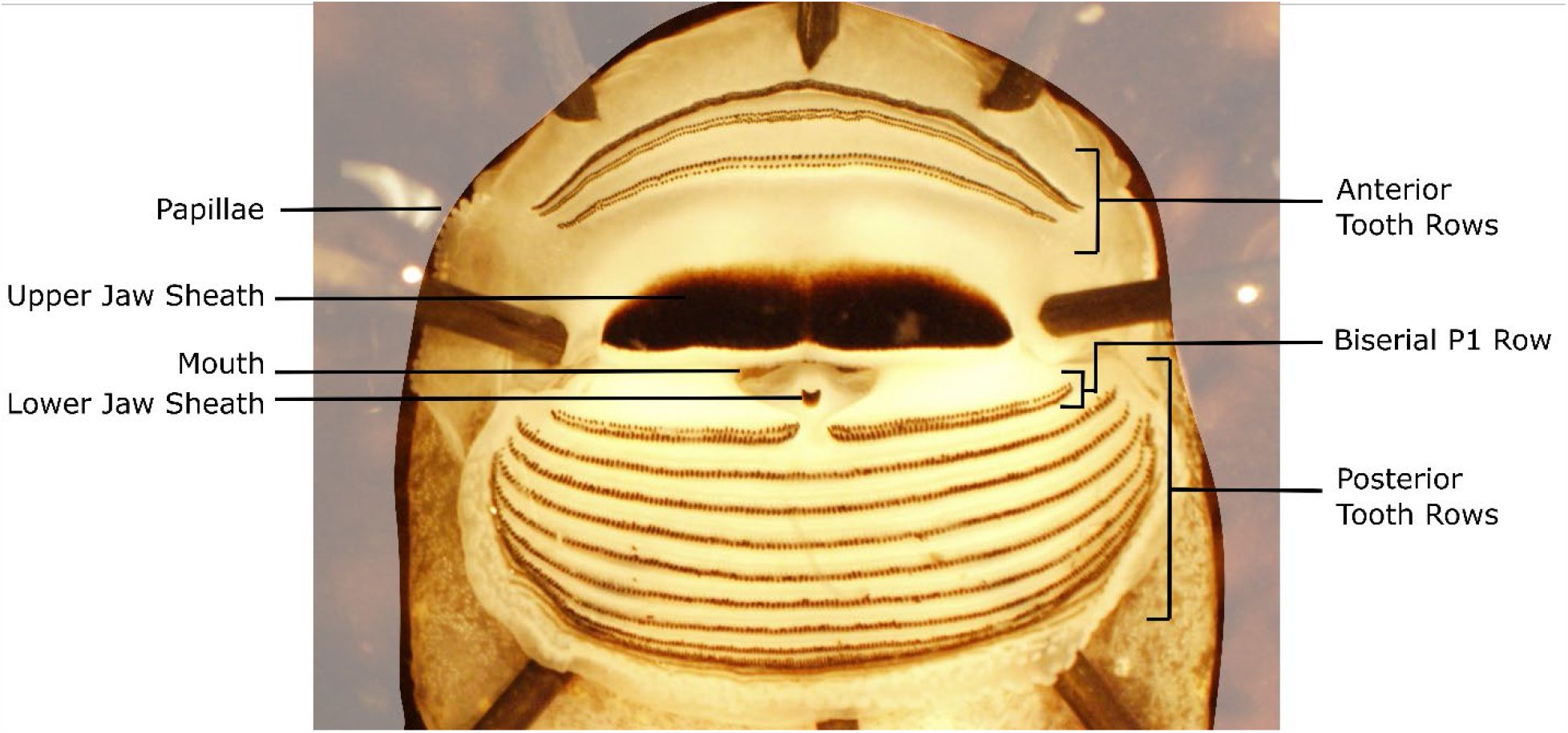
Pinned specimen of *Ascaphus montanus* (specimen ID WCF08228 from Lost Horse Creek, MT) showing labelled oral morphology, including labial tooth rows. Rows are numbered from anterior to posterior. The A2, A3, and P1 rows are biserial (i.e., have two rows of teeth), as demonstrated by the labelled P1 row. The labial tooth row formula (see Methods) for this specimen is 3/9(1), where (1) indicates the medial gap in the P1 row.

Tadpole oral morphology is simultaneously subject to developmental and extrinsic pressures. Individual rates of growth (cell differentiation) and development (cell specialization) may vary within and among populations and can be reflected in oral morphology (Altig and McDiarmid, 1999; Conradie and Conradie, 2015). Temperature influences rates of growth and development (Gillooly et al., 2002; Angilletta Jr and Dunham, 2003; Angilletta et al., 2004; Gomez-Mestre et al., 2010; Liess et al., 2013), and may shape oral morphology through this interaction with developmental processes. For example, Vences et al. (2002) demonstrated the downstream effects of temperature-driven developmental plasticity on oral morphology in tadpoles of the common frog, *Rana temporaria*, as temperature effects on developmental rates led to variation in the number of labial teeth and of labial tooth rows. Additionally, temperature plays a large role in many other ectothermic processes, such as metabolic rate (Gillooly et al., 2001; Brown et al., 2004) as well as rates of feeding (De Sousa et al., 2015), demonstrating the potential to influence oral morphology independent of development. Therefore, to uncover patterns in tadpole oral morphology, both the independent and interacting effects of development and temperature must be considered.

Tailed frogs (Family: Ascaphidae) occupy natural temperature gradients along both latitude and elevation, providing an opportunity to investigate these drivers within species. The two species in this family occupy cold, fast-flowing, high gradient streams in mesic, forested landscapes of the Klamath Mountains, Coast Ranges, and Cascade Mountains (*Ascaphus truei*), and northern Rocky Mountains (*A. montanus*) of the United States and Canada (Figure 2). Tailed frog larval development is slow (Brown, 1975; 1989) and variable, lasting one to four years, with greater variation found across the wider latitudinal and elevational range of *A. truei* (Hayes and Quinn, 2015). This variation in developmental rates may be affected by environmental conditions (Brown, 1990; Wallace and Diller, 1998; Bury and Adams, 1999). Tailed frog tadpoles also have a distinctive oral morphology with specialized, suctorial mouthparts. The tadpoles have an enlarged oral disc with a sucker and flexible papillae that attach to stream substrate and prevent them from being washed downstream (Gradwell, 1971). Unlike many frog species, labial teeth of *Ascaphus* tadpoles are thought to have a reduced role in tadpole stabilization (Gradwell, 1971). Instead, labial teeth in these species largely function in feeding kinematics by lifting food off of rock substrates (along with the jaw sheaths) and moving it to the mouth (Gradwell, 1971; Altig and Johnston, 1989). The well-understood function of labial teeth, varying developmental rates, and a range that traverses many ecological gradients make tailed frogs an excellent system to investigate the drivers of variation in labial tooth number, a trait that has not yet been investigated in detail across multiple populations of anuran tadpoles.

**Figure 2.**
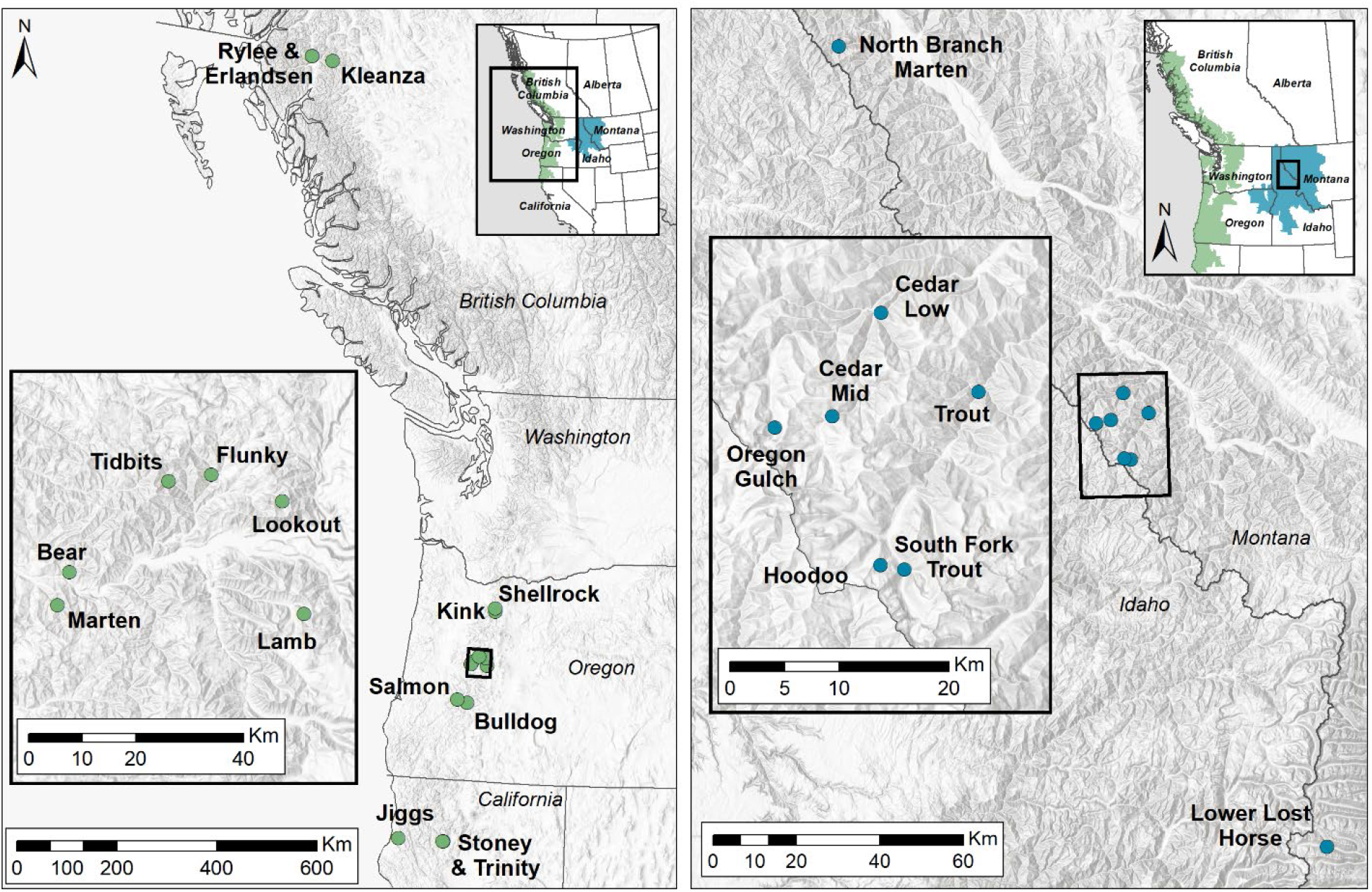
Sampling locations for *Ascaphus truei* (left) and *Ascaphus montanus* (right). Inset maps show estimated range boundaries for *A. truei* (in green) and *A. montanus* (in blue). Range maps from NatureServe & IUCN (2012).

Here, we investigated patterns and drivers of variation in tadpole labial tooth number among populations of both tailed frog species. By sampling populations spanning both elevation (both species) and latitudinal (*A. truei*) gradients, we tested the roles of development, local habitat temperature, and their interaction on shaping these patterns. We counted multiple tooth rows in tadpoles from each sampled population, in addition to measuring length and assessing developmental stage. To characterize local thermal conditions, we recorded stream temperatures from each sampled site. We expected variation in development and local temperature conditions to influence variation in labial tooth number within species. Tadpoles inhabiting warmer streams may require more energy to meet metabolic demands (Gillooly et al., 2001; Liess et al., 2013) and more labial teeth may be more efficient for feeding (Venesky et al., 2010a; b). We therefore expect populations from warmer streams to have more labial teeth than those occupying colder streams in response to environmental pressures for increased feeding efficiency. Alternatively, populations from colder streams may have lower developmental rates as a response to temperature (i.e., developmental plasticity). If this relationship is driving labial tooth number variation, we predicted that colder populations would have more labial teeth than those from warmer streams as an outcome of greater energy allocation to growth.

## MATERIALS AND METHODS

### Tadpole collection

At each site we sampled stream reaches (∼100 m in length) in the summers of 2016 and 2017 across British Columbia (N=3), Oregon (N=10), California (N=3), and Montana (N=8; Figure 2; site coordinates available in Supplemental Table A). We focused sampling within ∼100m stream reaches as *Ascaphus* tadpoles are not highly vagile (Gaige, 1920; Altig and Brodie, 1972), having been observed moving a mean total 1.1m/day in natural environments similar to the streams we sampled (Wahbe and Bunnell, 2001). We captured tadpoles by shifting and brushing rocks and benthic substrate while holding an aquarium net downstream. Tadpoles were held in coolers in the field, and were later euthanized, photographed laterally on a grid, fixed in formalin, and stored in 95% ethanol. All tadpole collection and euthanasia were performed under the following permits: California Department of Fish and Wildlife permit number SC-12131; Oregon Department of Fish and Wildlife permit numbers 117-16 and 110-17; Montana Department of Fish, Wildlife, and Parks permit numbers 2016-100 and 2017-060; Simon Fraser University Animal Care Committee protocol 1130B-14; and Institutional Animal Care and Use Committee protocol 16-6667AA.

### Labial teeth data collection

For each sampled site, we chose ten specimens (160 *A. truei* and 80 *A. montanus*) for labial tooth counts. We selected specimens that were large enough to pin and photograph without destroying the specimen (Gosner stage > 28; Gosner 1960), and for which the developmental stage was not so advanced that the suctorial mouth was reduced (Gosner stage < 41). Specimens were pinned with the mouth open to show a full view of the labial teeth rows. We took at least three photographs of each specimen using an Olympus SZX10 scope: full view of mouth, anterior rows only, and posterior rows only (Supplemental Figure A). All counts were done by one author (CMM).

For each specimen, we first calculated the labial tooth row formula (LTRF), designated as the number of rows in the anterior labium (A) / the number of rows in the posterior labium (P). Anterior and posterior rows were numbered from anterior to posterior within the labium (Altig and McDiarmid, 1999). The most posterior row was defined as requiring distinct, discernible teeth throughout the row. If upon zooming in on a row, no distinct teeth were discernible, the row was not included in the LTRF. A and P values were followed by a row number in parentheses if a medial gap was present in the row. For example, a tadpole with 3 anterior rows and 7 posterior rows would have an LTRF of 3/7. If the first posterior row of teeth had a medial gap, then the LTRF would be 3/7(1). Rows consisting of two sub-rows on each tooth ridge were recorded as biserial and referred to as anterior or posterior.

We then used Adobe Photoshop and ImageJ (Rasband, 2018) to count individual teeth in the following rows: A3 anterior, A3 posterior, P1 posterior, P2, and P3. In specimens where the P2 row was biserial, we counted the P2 posterior row. These five rows were selected because their ends were consistently discernible from the center to the outer edges of the oral disc. Each tooth was individually marked with a red dot in ImageJ and counted moving from left to right. The same count was then performed with a dot of a different color moving from right to left. If the two counts matched, the number of teeth was recorded for the row. If the two counts were not the same, a third count was performed to serve as a tiebreaker. In a small number of cases, counts were initially listed as “NA” for one of the following reasons: a photograph was too blurry for all teeth to be discernible, there were teeth missing, there was damage to the tooth row, or the view of the teeth was obstructed. In these cases, we filled in NA values using one or more of the following approaches: (1) we used ImageJ to sharpen blurry edges to improve count accuracy; (2) we re-photographed the specimen; and/or (3) we estimated tooth counts based on the spacing in the rest of the row in areas of glare or missing teeth.

We counted individual teeth in all five rows for 114 individuals. Based on correlations in counts across the five rows of 114 individuals, we reduced the number of rows counted to A3 anterior, A3 posterior, and P2 for the remaining 126 specimens (total N=240). Rows P1 and P3 were not analyzed since they were highly correlated with P2 (r > 0.95 in both cases). P2 was retained over P1 because it was not biserial; P2 was retained over P3 because the P2 row photo quality was generally higher than P3 (e.g., depth of field was superior/least blurry at the edges of the oral disc).

### Staging and length measurements

Each tadpole was assigned a developmental stage based on a slightly modified staging system developed by Gosner (1960). The unique and slow development of *Ascaphus* tadpoles (Brown, 1975; 1990) required that we accommodate distinctive developmental features in *Ascaphus*, such as the late reduction of the suctorial disc and oral labia (manuscript authors, unpublished data). We used ImageJ to measure the length of each tadpole (tip of snout to tip of tail) from the lateral photographs; each tadpole was measured twice, and the values were averaged.

### Temperature predictor variables

Annual temperatures were calculated from continuous temperature loggers (Onset Hobo Pendants UA-001-064) installed at each site. Data loggers were installed at the upstream and downstream ends of the stream reach using rebar pounded into the substrate. Data loggers were protected from debris and sunlight using PVC housing drilled with holes to allow water flow. Temperature was recorded every 4 hours. Annual temperature for most sites was calculated on either a 2016-2017 or 2017-2018 water year (October 1 - September 30), although some sites used a different time frame or were missing small amounts of data (see Supplemental Table A for dates and loggers used in calculations for each site).

### Analyses

All analyses were performed using R v. 4.1.2 (R Core Team, 2021). We first determined if we could treat Gosner stage as continuous in our models, rather than as an ordered factor. For each of the three labial tooth rows, we ran two GLMs treating Gosner stage as either a continuous or ordinal variable. For these and all subsequent GLM models, we checked the goodness of fit of the GLM with a Poisson error distribution using a chi-square test based on the residual deviance and degrees of freedom. If the Poisson distribution did not fit the data, we used a negative binomial distribution fit with the R package *MASS* (Venables and Ripley, 2002). We then compared the models using a likelihood ratio test, AIC, and BIC. We ran this analysis separately for each species on each labial tooth row.

We described patterns in LTRF and tooth number variation. We tested for species differences in labial tooth counts using a GLM negative binomial distribution (based on goodness of fit test). We tested for within-species population variation in tooth counts using an ANOVA and tested our predictions regarding development and temperature using species-specific GLMs (α = 0.05) and calculated a pseudo-R^2^ for each model (Zuur et al., 2009). Tadpole size and developmental stage were highly correlated in both species (*A. montanus*: Pearson’s correlation coefficient = 0.91, t=20.01, df= 78, p<0.0001; *A. truei*: Pearson’s correlation coefficient = 0.62, t=9.81, df= 158, p<0.0001). Therefore, our models included tadpole developmental stage, stream average temperature, and the interaction between stream average temperature and developmental stage.

## RESULTS

### Gosner stage analyses

In both species, a Poisson GLM fit the A3 anterior and P2 data, while a negative binomial GLM was the best fit for the A3 posterior data. For the A3 anterior data, treating Gosner stage as a continuous variable was the better model according to AIC and BIC, with the likelihood ratio test showing no significant difference between the models. For the A3 posterior row, treating Gosner stage as a continuous variable was the better model according to BIC, with strong support for both models based on AIC. The likelihood ratio tests for A3 posterior showed a significant difference between the models for both *A. truei* (*p* = 0.01) and *A. montanus* (*p* = 0.04). In both species, the P2 data had the least support for a continuous treatment, with BICs in support of continuous treatment, but AIC and likelihood ratio tests with little to no support (full results in Supplemental Table B). Given these mixed results, we chose to treat Gosner stage as continuous for the remainder of analyses to simplify model fitting and interpretation.

### Patterns and drivers of labial teeth variation

All specimens had identical LTRF values for the number of anterior rows (3) but varied in the number of posterior rows (7-9). All specimens had a medial gap in the P1 row and were biserial in rows A2, A3, and P1. All 30 individuals from the three British Columbia sites also had a biserial P2 row (Supplemental Figure A), which was not observed in other specimens.

Final tooth counts for the three rows (A3 anterior, A3 posterior, and P2) were highly correlated (Pearson’s correlation coefficient > 0.74, Supplemental Table C). Therefore, we present the model results for the P2 row in the main text, as this row had the most amount of variation explained (A3 row results are provided in Supplemental Table E). Using a GLM with a negative binomial distribution, we found that *A. truei* had higher P2 tooth counts than *A. montanus* (Supplemental Table D) and that populations of both species varied in their P2 counts (Figure 3; *A. montanus* ANOVA F _(7,72)_=27.39, p<0.0001; *A. truei* ANOVA F_(15,144)_=12.60, p<0.0001).

**Figure 3.**
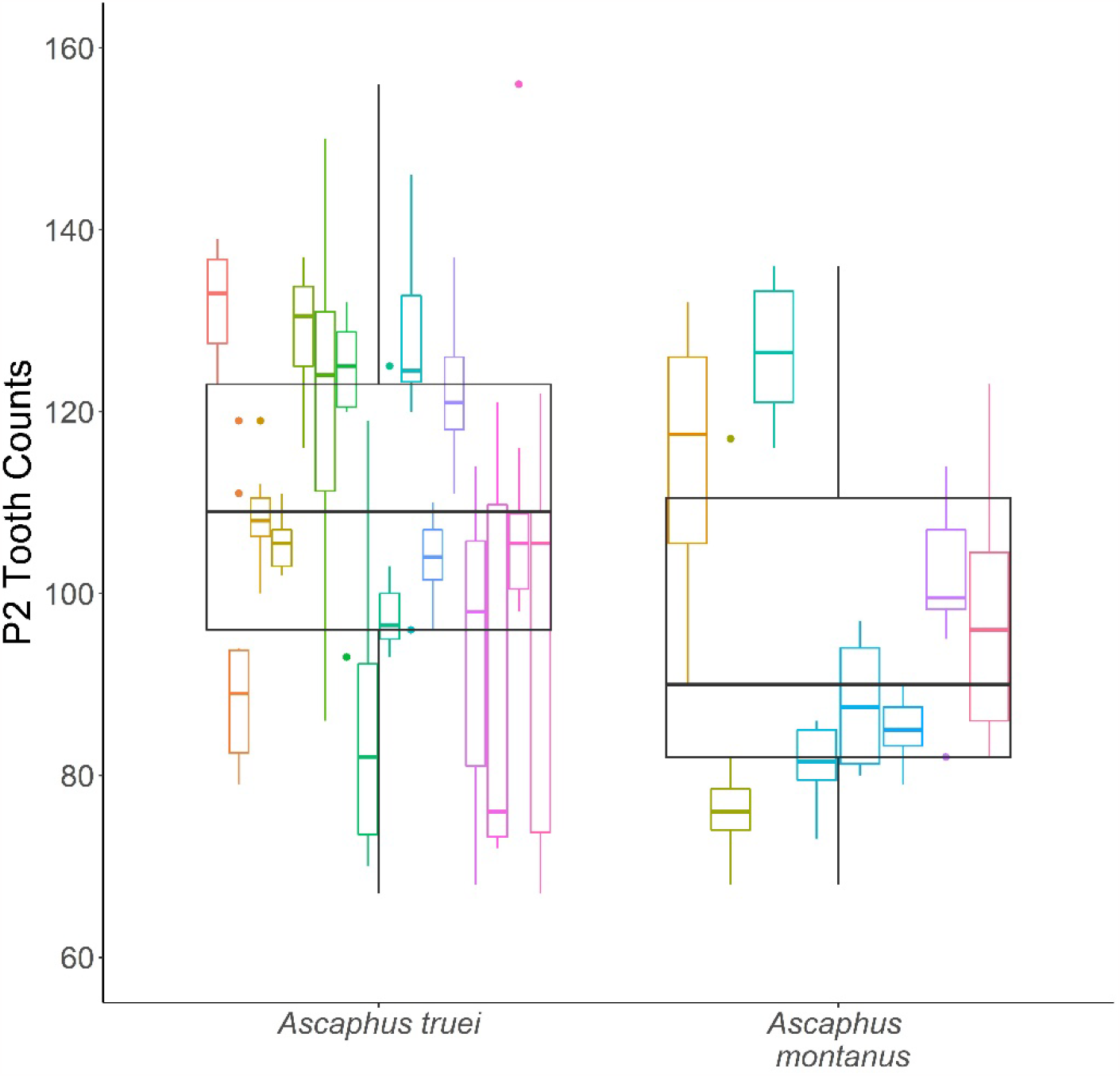
*Ascaphus truei* tadpoles had higher overall counts of labial teeth in the P2 row than *A. montanus* tadpoles. Within both species, populations (here shown as different colored boxplots) varied in their P2 tooth counts.

Developmental stage significantly affected tooth counts positively in both species. Stream temperature had a significant, positive effect on tooth counts in *A. truei* only, such that tadpoles from warmer streams had more teeth. The interaction of these two predictors trended positively in *A. montanus*, such that the relationship between developmental stage and tooth number increased in warmer streams. In *A. truei*, however, the interaction term was significantly negative, such that the relationship between developmental stage and tooth counts decreased in warmer streams (Figure 4). The model for *A. montanus* explained 67% of the variation in labial tooth number; the model for *A. truei* explained 77% of variation in labial tooth number. Despite the high correlations among tooth rows, there were slight differences in model results for the other tooth rows: Temperature was a significant, positive predictor of tooth number in the *A. montanus* models for A3 anterior and posterior rows, and the interaction of the two predictors was not a significant predictor of the A3 posterior tooth counts in *A. truei* (Supplemental Table E).

**Table 1.** Results from species-specific GLMs investigating the drivers of A3 anterior labial tooth number variation.

|  | Predictor | Estimate (SE) | Z-value | p-value |
| --- | --- | --- | --- | --- |
| *Ascaphus truei* | Intercept | 4.68 0.01 | 572.58 | < 0.0001 |
|  | Developmental Stage | 0.14 0.01 | 16.69 | < 0.0001 |
|  | Average Temperature | 0.05 0.01 | 5.07 | < 0.0001 |
|  | Developmental Stage: Average Temperature | -0.03 0.01 | -2.78 | 0.005 |
| *Ascaphus montanus* | Intercept | 4.55 0.01 | 355.39 | < 0.0001 |
|  | Developmental Stage | 0.15 0.01 | 11.48 | < 0.0001 |
|  | Average Temperature | 0.02 0.01 | 1.42 | 0.15 |
|  | Developmental Stage: Average Temperature | 0.02 0.01 | 1.84 | 0.066 |

**Figure 4.**
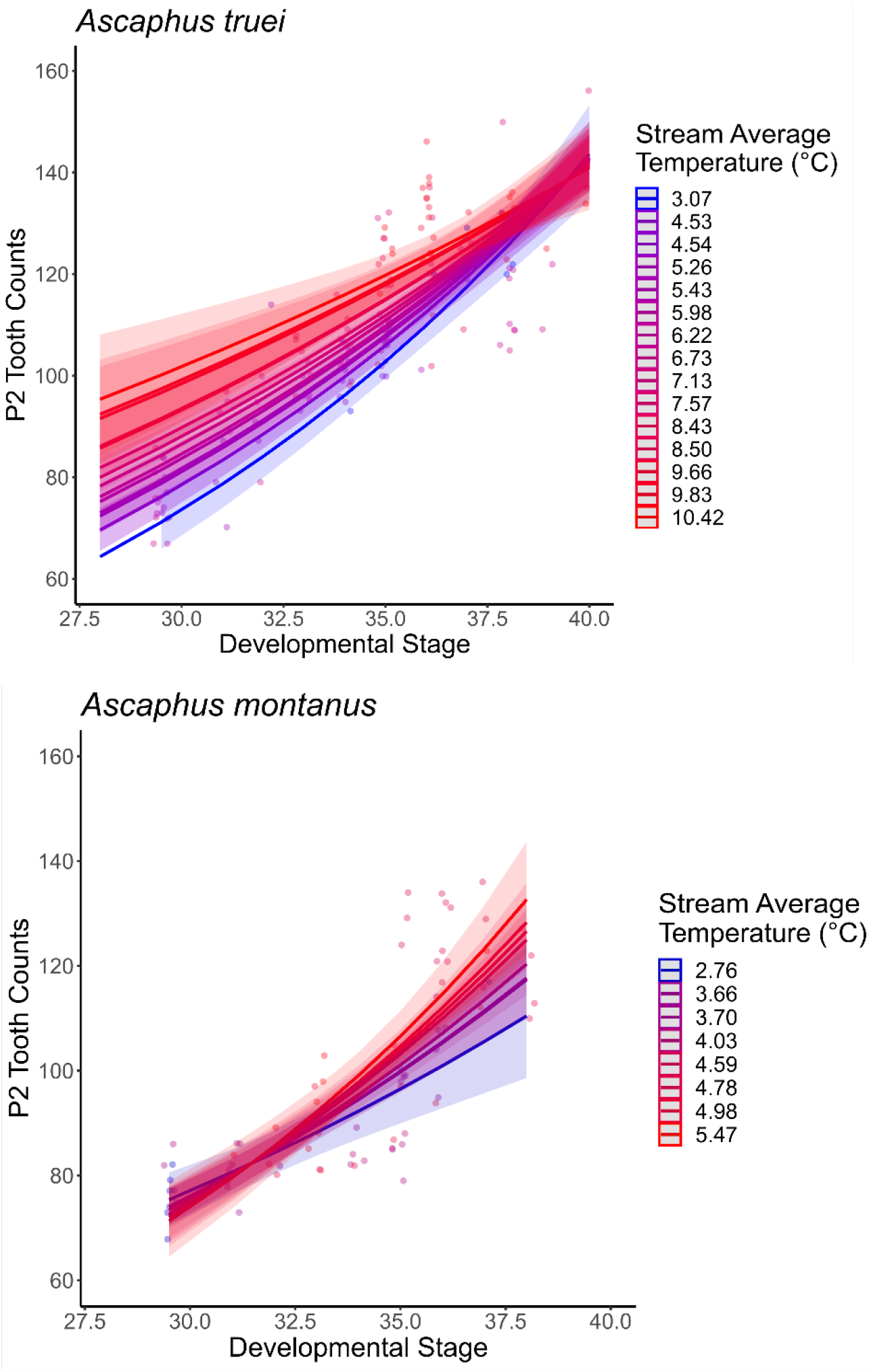
From the results of the species-specific GLMs, we predicted relationships (shown with 1.96*SE confidence intervals) between P2 tooth counts and developmental stage at each temperature point sampled (annual average stream temperature). Raw data points are also shown on these plots.

## DISCUSSION

Tadpole oral morphology has played an important role in clarifying relationships among diverse species of frogs (Altig, 2006; Vera Candioti and Altig, 2010). Despite this, few studies have investigated patterns of intraspecific variation in oral morphology. Using a cold-water specialist frog system (*Ascaphus spp*.), we quantified variation in the numbers of labial teeth among populations of both tailed frog species. As expected, we identified interconnected relationships between tadpole development and local stream temperature, explaining up to 77% of the variation in labial tooth number. We found that labial tooth number variation is independently influenced by development and temperature, as well as their interaction (i.e., developmental plasticity). We also uncovered patterns in labial tooth rows, including a bifurcation of the P2 tooth row unique to *A. truei* tadpoles from northern British Columbia, Canada. Broadly, these results offer insights into the evolution of an understudied oral morphological trait in tadpoles and emphasize the interrelated roles of development and local environment in shaping larval traits.

### Drivers of labial tooth number variation

Development is a key predictor of tadpole oral morphological traits (Thibaudeau and Altig, 1988; Altig and McDiarmid, 1999). As expected, developmental stage was positively related to labial tooth numbers among populations of both species of tailed frogs, such that tadpoles gained teeth throughout development. Developmental stage was also related to tadpole size in both species, although more strongly in *A. montanus*.

The stronger relationship in *A. montanus* may be due to differences in sample sizes, though may also reflect differences between growth and development between the species. Nevertheless, these relationships suggest that tadpoles are experiencing allometric growth during development as larger, more developed tadpoles would have larger bodies, which could support more teeth.

Similarly, Conradie and Conradie (2015) found positive relationships between tadpole size and the number of posterior labial tooth rows in four species of ghost frogs (family: Heleophrynidae), demonstrating an increase in number of overall labial teeth with body size. Beyond the independent effects of developmental stage on labial tooth number variation, our results demonstrated that temperature effects on development also play a role.

Growth and development in tadpoles are plastic traits and can be influenced by environmental variables such as temperature (Newman, 1992; Denver et al., 1998; Ruthsatz et al., 2018). For example, warm temperatures may prompt a shift in energy allocation towards development rather than growth as high temperatures are associated with increased metabolic demand and rates of development (Gillooly et al., 2002; Angilletta Jr and Dunham, 2003; Angilletta et al., 2004; Gomez-Mestre et al., 2010; Liess et al., 2013). Our results support temperature as a cue for developmental plasticity resulting in variation in labial teeth number within both species. In *A. truei*, our data show that less-developed tadpoles vary in their labial tooth number across temperature but converge over development. In *A. montanus*, we observed the opposite trend, where tadpoles developing in warmer streams gain more teeth throughout development than those in colder streams. The underlying mechanism driving these relationships requires further investigation.

Beyond the interacting effects of temperature on labial tooth number through development, our results demonstrate that warmer streams have a positive influence on labial tooth number. The energetic demands of occupying warmer streams may favor increased labial teeth number to improve feeding efficiency. De Sousa et al. (2015) found that tadpoles feeding in warm temperatures had faster gape cycles and a smaller maximum gape, suggesting a tradeoff between feeding speed and surface area fed upon. Additionally, Venesky et al. (2010a) investigated the effects of missing labial teeth on tadpole feeding kinematics and found that tadpoles with fewer labial teeth compensated for decreased feeding efficiency by increasing gape cycles. These two studies demonstrate that both increased temperatures and decreased labial tooth numbers may reduce food consumption without compensatory responses. Thus, maximizing the amount of food consumed in quick gape cycles may be particularly important in warm streams to maintain a large enough energy budget to support development and growth. Similarly, increased numbers of labial teeth may be evolved responses to increase feeding efficiency in warmer temperatures.

The relationships we uncovered may also be related to factors not investigated in this study. For example, the observed increase in labial teeth number at higher stream temperatures may be partly driven by increased predation pressure. In these habitats, fish and crayfish predators are more abundant at lower elevations where temperatures are warmer (Reeves et al., 1998; Creed, 2006). *Ascaphus* tadpoles alter their feeding behavior in the presence of predator cues, reducing time spent feeding during the day (Feminella and Hawkins, 1994). Increased feeding efficiency through increased number of labial teeth may thus reflect a response to compensate for decreased feeding time. Food availability may also be contributing to these patterns. Canopy cover among sampled streams may vary, influencing the amount of direct sunlight on the stream and thus primary productivity (Hawkins et al., 1983). Areas with less canopy cover may have increased food availability for tadpoles, which would lead to higher rates of growth (including increased number of labial teeth) and development (Kiffney and Richardson, 2001). Both increases in temperature and changes in canopy cover are associated with decreases in *Ascaphus* populations (Hossack et al., 2023). Therefore, further studies investigating the relationships we uncovered between number of labial teeth and temperature, as well as testing additional hypotheses regarding predation risk and resource availability, are necessary to understand how changes to the environment will ultimately impact populations of tailed frogs.

### Broad patterns in labial tooth variation

Despite the broad geographic sampling across the two species, the labial tooth row formula (LTRF) was relatively consistent, with variation only in the number of posterior rows. This variation is likely due to developmental stage and size, as has been shown in other species (Bresler and Bragg, 1954; Conradie and Conradie, 2015).

Interestingly, all 30 tadpoles from northern British Columbia had a biserial P2 row, which was not found in any of the other 210 *Ascaphus* individuals. Bifurcation of the tooth row may be related to increased flexibility (Annibale et al., 2020), suggesting a possible functional role, and/or differences in developmental sequences (Vera Candioti et al., 2011; Grosso et al., 2019). Further investigation is needed to determine the geographic extent and underlying cause of the unique P2 bifurcation.

We also found that *Ascaphus truei* had more teeth than *A. montanus*. These species differences in overall tooth counts may reflect evolutionary history, the outcome of neutral processes since divergence, or ecological differences. *A. truei* and *A. montanus* diverged from their common ancestor over 5 million years ago, following the rise of the Cascade Mountain range in North America and associated climatic changes (Nielson et al., 2001). Despite both occupying montane streams, these species experience different environmental conditions that reflect their coastal (*A. truei*) and continental (*A. montanus*) climate regimes. From the sites we sampled for the present study, we found that *A. truei* streams had higher average temperatures than *A. montanus*, as well as greater variation in average temperature. As temperature was a positive driver of tooth variation in all tooth rows for both species, higher average temperatures in *A. truei* may explain why this species has more teeth overall. However, this study did not seek to understand trait divergence between the species and further investigation into the influences of ecological differences and evolutionary history is needed.

## Acknowledgements

For assistance with data collection and field logistics, we thank A.A. Shah, D. Oliver, L. King, A. Breda, J. Suh, K. Pain, R. Gimple, J. Kendrick, R. Jackson, R. Murray, W. Palen, B. Hossack, and W. Lowe. We also thank the Devo-Evo discussion group led by Kim Hoke, Rachel Mueller, and Ashok Prassad for thoughtful discussions regarding development and evolution. We acknowledge funding from a National Science Foundation (NSF) Rules of Life grant (1838282) to W. Chris Funk and a Natural Sciences and Engineering Research Council of Canada (NSERC) post-graduate scholarship to Amanda S. Cicchino (PGSD2-532408-2019).

## Data Accessibility

Supplemental material is available at https://www.ichthyologyandherpetology.org/XXX.

## Supplementary Material

**Supplemental Figure A.**
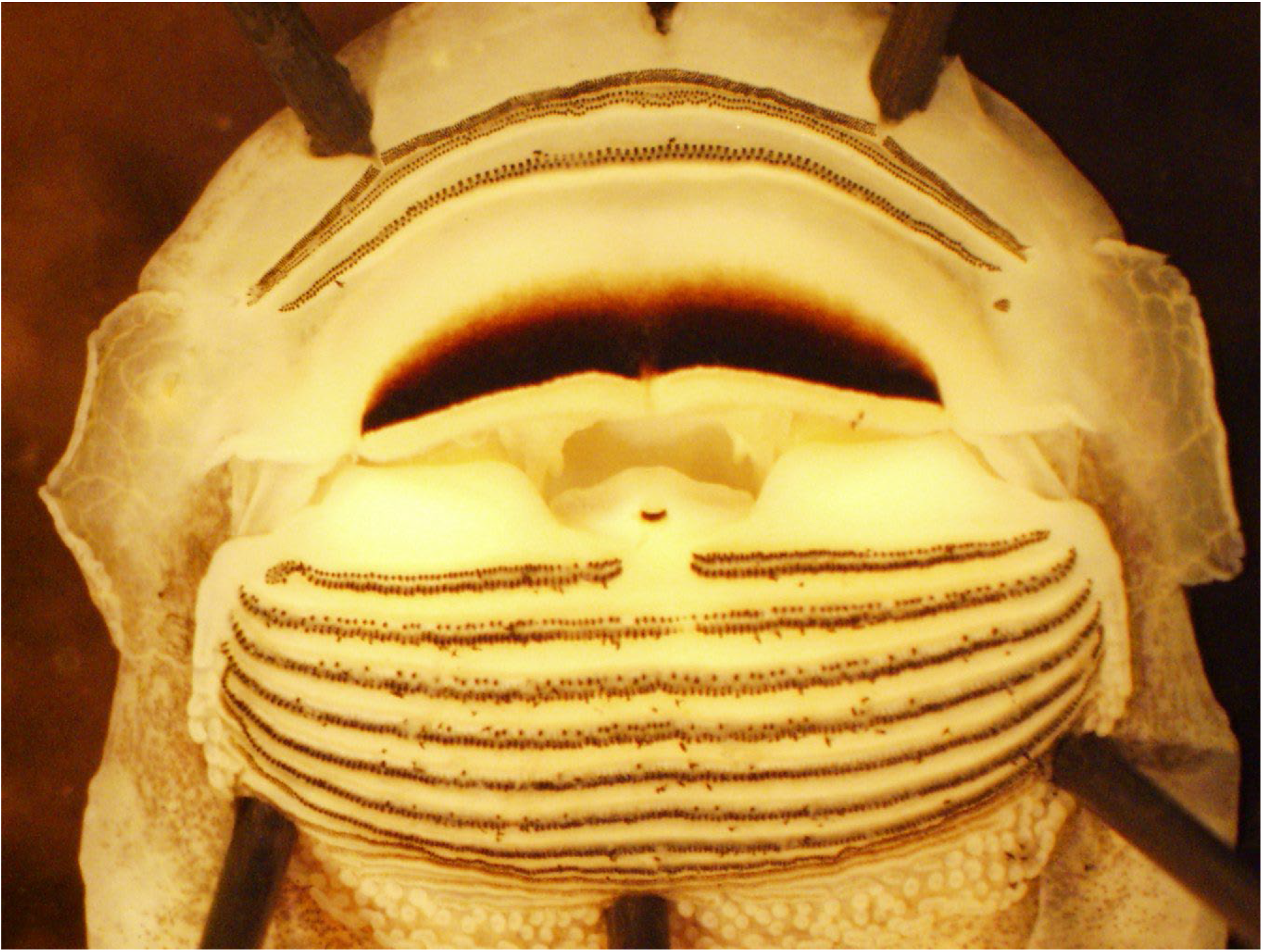
Pinned specimen of *Ascaphus truei* (specimen ID WCF07243) from Kleanza Creek, BC. The 30 tadpoles collected from BC all had a biserial P2 row, shown here, which was not observed in other *A. truei* tadpoles.

**Supplemental Table A.**
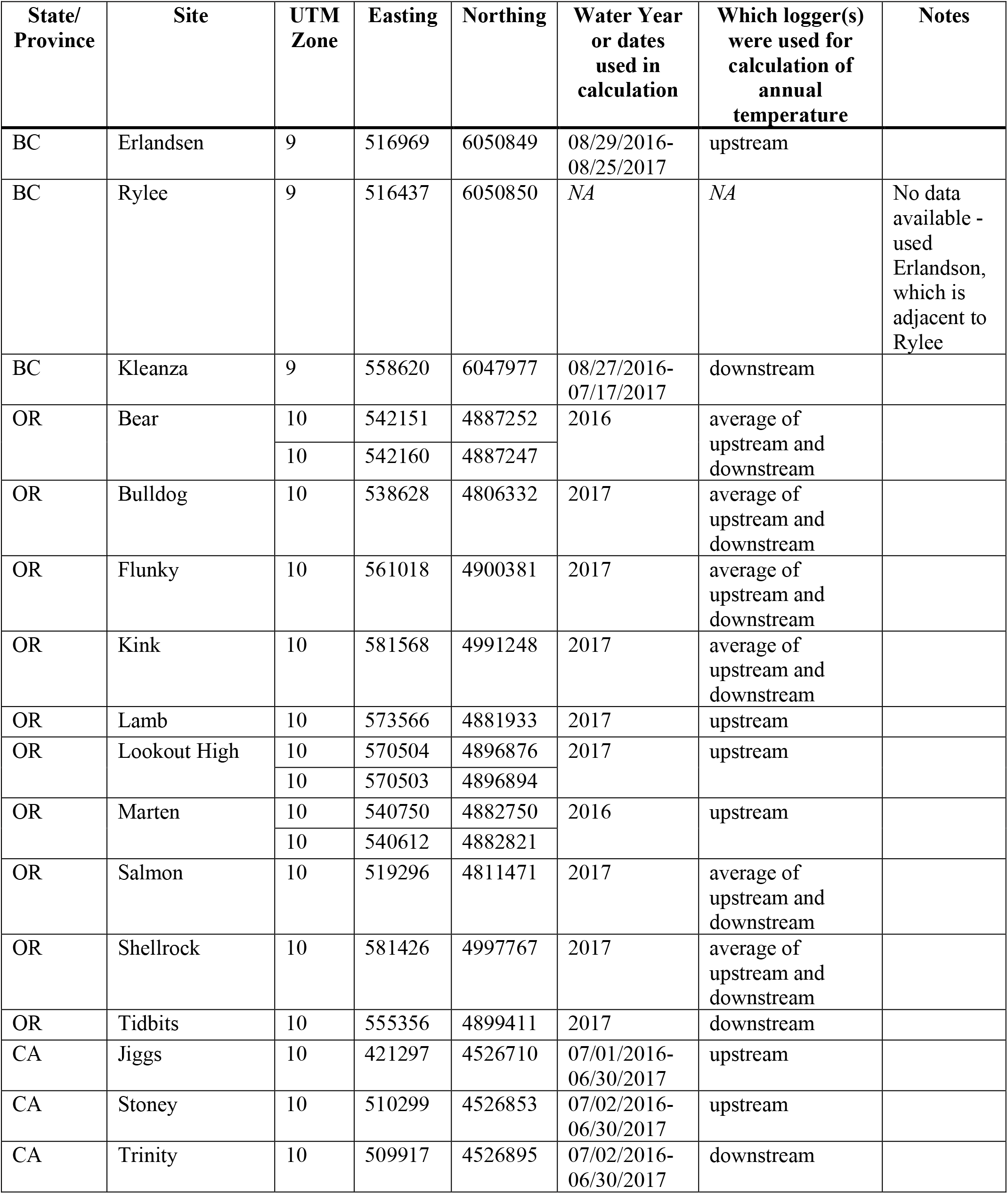
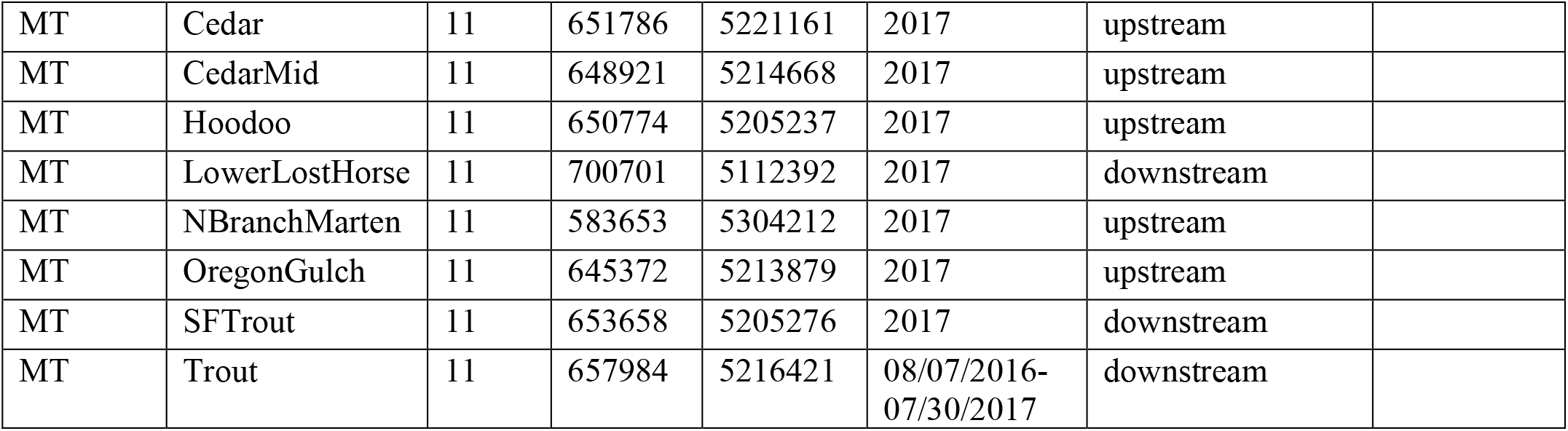
Temperature logger metadata.

**Supplemental Table B.** AIC, BIC, and likelihood ratio test results for GLM models of tooth count by Gosner stage, fit as continuous and ordinal.

| Species | Tooth Row | Model fit | AIC ordinal | AIC continuous | ΔAIC | BIC ordinal | BIC continuous | ΔBIC | LR p-value |
| --- | --- | --- | --- | --- | --- | --- | --- | --- | --- |
| *Ascaphus truei* | A3 anterior | poisson | 1183.6 | 1174.7 | 8.9 | 1220.5 | 1180.8 | 39.7 | 0.351 |
|  | A3 posterior | negative binomial | 1335.0 | 1337.2 | -2.2 | 1375.0 | 1346.4 | 28.6 | 0.014 |
|  | P2 | poisson | 1173.7 | 1199.4 | -25.7 | 1210.6 | 1205.6 | 5.0 | < 0.001 |
| *Ascaphus montanus* | A3 anterior | poisson | 605.4 | 601.3 | 4.1 | 626.9 | 606.0 | 20.9 | 0.197 |
|  | A3 posterior | negative binomial | 676.8 | 677.6 | -0.8 | 700.6 | 684.7 | 15.9 | 0.039 |
|  | P2 | poisson | 603.9 | 608.5 | -4.6 | 625.3 | 613.3 | 12.0 | 0.009 |

**Supplemental Table C.** Correlations among final tooth row counts (N=240) were highly correlated.

|  | Tooth Rows | Pearson's correlation coefficient | t-value, df | p-value |
| --- | --- | --- | --- | --- |
| *Ascaphus montanus* | A3 anterior – A3 posterior | 0.89 | 17.39, 78 | $2.2e^{-16}$ |
| | A3 anterior – P2 | 0.87 | 15.46, 78 | $2.2e^{-16}$ |
| | A3 posterior – P2 | 0.80 | 11.73, 78 | $2.2e^{-16}$ |
| *Ascaphus truei* | A3 anterior – A3 posterior | 0.87 | 22.28, 158 | $2.2e^{-16}$ |
| | A3 anterior – P2 | 0.85 | 20.18, 158 | $2.2e^{-16}$ |
| | A3 posterior – P2 | 0.74 | 13.90, 158 | $2.2e^{-16}$ |

**Supplemental Table D.** Results from GLM models testing for species differences in tooth number for each investigated tooth row.

| Tooth Row | Model fit | Parameter (*A. montanus* as reference factor level) | Fixed effect estimate | Fixed effect std. error | Fixed effect z-value | Fixed effect p-value | Pseudo-R<sup>2</sup> |
| --- | --- | --- | --- | --- | --- | --- | --- |
| P2 | negative binomial | Intercept | 4.57 | 0.02 | 220.85 | <0.0001 | 0.80 |
|  |  | *A. truei* | 0.11 | 0.02 | 4.59 | <0.0001 |  |

**Supplemental Table E.** Results from GLM models testing drivers in the A3 anterior (A3a) and posterior (A3p) tooth rows. Despite these tooth rows being highly correlated with the P2 counts, there were slight differences in the relationships with predictors.

| A3a (Pseudo-R<sup>2</sup>) | Predictor | Estimate (SE) | Z-value | p-value |
| --- | --- | --- | --- | --- |
| *Ascaphus truei*<br>(67%) | Intercept | 4.70 (0.01) | 581.13 | < 0.0001 |
|  | Developmental Stage | 0.09 (0.01) | 10.99 | < 0.0001 |
|  | Average Temperature | 0.04 (0.01) | 4.30 | < 0.0001 |
|  | Developmental Stage:<br>Average Temperature | -0.03 (0.01) | -2.59 | 0.01 |
| *Ascaphus montanus*<br>(60%) | Intercept | 4.57 (0.01) | 361.77 | < 0.0001 |
|  | Developmental Stage | 0.11 (0.01) | 0.85 | < 0.0001 |
|  | Average Temperature | 0.03 (0.01) | 2.42 | 0.01 |
|  | Developmental Stage:<br>Average Temperature | 0.02 (0.01) | 2.03 | 0.04 |

| A3P (Pseudo-R<sup>2</sup>) | Predictor | Estimate (SE) | Z-value | p-value |
| --- | --- | --- | --- | --- |
| *Ascaphus truei*<br>(54%) | Intercept | 4.75 0.01 | 438.65 | < 0.0001 |
|  | Developmental Stage | 0.11 0.01 | 10.17 | < 0.0001 |
|  | Average Temperature | 0.05 0.01 | 4.23 | < 0.0001 |
|  | Developmental Stage:<br>Average Temperature | -0.02 0.01 | -1.29 | 0.20 |
| *Ascaphus montanus*<br>(58%) | Intercept | 4.60(0.02) | 240.67 | < 0.0001 |
|  | Developmental Stage | 0.17 0.02 | 8.27 | < 0.0001 |
|  | Average Temperature | 0.05 0.02 | 2.45 | 0.01 |
|  | Developmental Stage:<br>Average Temperature | 0.04 0.02 | 2.40 | 0.02 |

## Notes

### Competing Interest Statement

The authors have declared no competing interest.

### Summary of Updates

Manuscript reframed.

